# Polygenic pathogen networks control of host and pathogen transcriptional plasticity in the Arabidopsis-Botrytis pathosystem

**DOI:** 10.1101/2023.03.16.533032

**Authors:** Parvathy Krishnan, Celine Caseys, Nik Soltis, Wei Zhang, Meike Burow, Daniel J. Kliebenstein

## Abstract

Bidirectional flow of information shapes the outcome of the host-pathogen interactions and depends on the genetics of each organism. Recent work has begun to use co-transcriptomic studies to shed light on this bidirectional flow, but it is unclear how plastic the co-transcriptome is in response to genetic variation in both the host and pathogen. To study co-transcriptome plasticity, we conducted transcriptomics using natural genetic variation in the pathogen, Botrytis cinerea, and large effect genetic variation abolishing defense signaling pathways within the host, Arabidopsis thaliana. We show that genetic variation in the pathogen has a greater influence on the co-transcriptome than mutations that abolish defense signaling pathways in the host. Genome wide association mapping using the pathogens genetic variation and both organisms’ transcriptomes allowed an assessment of how the pathogen modulates plasticity in response to the host. This showed that the differences in both organism’s responses were linked to trans-eQTL hotspots within the pathogen’s genome. These hotspots control gene sets in either the host or pathogen and show differential allele sensitivity to the hosts genetic variation rather than qualitative host specificity. Interestingly, nearly all the trans-eQTL hotspots were unique to the host or pathogen transcriptomes. In this system of differential plasticity, the pathogen mediates the shift in the co-transcriptome more than the host.

## Introduction

How hosts and microbes interact depends on a massive and rapid flow of information between the organisms (Kang 2019). For one organism to effectively shift the interaction, this information flow has to be received and transformed into appropriate responses, such as the coordinated and orchestrated action of innumerable signaling molecules, regulatory cascades and metabolic pathways (Botero *et al*. 2018; Ma *et al*. 2021). In a successful interaction, the organism(s) responses are sustainable in their ever-changing complex micro and macro environment encompassing a range of symbiotic and pathogenic organisms also engaging in cross-species communication (Weiland-Bräuer 2021). Overall, the flow of information is shaped at the molecular level into transcriptome, protein and/or metabolism responses (Szymański *et al*. 2020; Chen *et al*. 2021). Understanding the information flows between interacting organisms is essential to characterize the underlying biological processes that lead to differential phenotypic outcomes ranging from disease to beneficial symbiotic interactions.

Studies of plant-pathogen interactions often focus on the flow of information mediated by a myriad of effector molecules from the pathogen to the host (Bent and Mackey 2007; Boller and Felix 2009) with a response by the host generally following the gene for gene interaction model (HH, Flor 1942). This includes an array of small secreted effector proteins, hydrolysis enzymes like plant cell wall degrading enzymes, oligosaccharides, specialized metabolites and small RNAs (Weiberg *et al*. 2013; Wang *et al*. 2016; van der Does and Rep 2017; Quoc and Bao Chau 2017). Plants have in turn evolved the ability to interpret these pathogen signals, and mount defense responses by combining various signal transduction mechanisms, including mitogen-activated kinases (MAPK), reactive oxygen species (ROS), and phytohormones like jasmonic acid (JA), ethylene (ET), and salicylic acid (SA) pathways in addition to their crosstalk. The end-point of these signal cascades is frequently the production and or transport of specialized metabolites like glucosinolates, camalexin, terpenes, alkaloids and phenylpropanoids that can poison the pathogen (Rogers 1996; Sticher *et al*. 1997; Bednarek *et al*. 2009; Stotz *et al*. 2011; Shlezinger *et al*. 2011; Ahuja *et al*. 2012). Recent studies showed that these defense metabolites are then perceived by the pathogen and lead to corresponding changes in the attacking pathogens transcriptome indicating the presence of bidirectional information flow (Vela-Corcía *et al*. 2019; Kusch *et al*. 2022). This response/counter-response model in the host and pathogen transcriptomes suggests that it is possible to measure the bidirectional flow of information using a co-transcriptome approach, a simultaneous assessment of both transcriptomes.

In the last decade, co-transcriptomic studies have started to decipher the flow of information between interacting organisms (Kawahara *et al*. 2012; Yazawa *et al*. 2013; Jupe *et al*. 2013; Hacquard *et al*. 2013; Rudd *et al*. 2015; Wang *et al*. 2016; Dobon *et al*. 2016). A primary focus of these studies has been to query how qualitative effectors released from specialist plant pathogens with a limited host range lead to the transcriptional reprograming of the host transcriptome. For example, co-transcriptome studies of the rice blast fungus (Kawahara *et al*., 2012) and barley powdery mildew, *Blumeria graminis* f. sp. *hordei* (Bgh) (Hacquard *et al*. 2013) pathosystems showed infection-responsive expression patterns that diverge between compatible and incompatible interactions. The focus of these studies on systems with qualitative loci lead to a biallelic survey of genetic variation linked to presence/absence of these individual large-effect loci (Zhong *et al*. 2017; Yang *et al*. 2021).

In contrast to qualitative systems, most plant-pathogen interactions are not guided by large-effect loci. For example, plant interactions with generalist necrotrophic pathogens like *Botrytis cinerea* and *Sclerotinia sclerotiorum* are shaped by a myriad of moderate to small effect loci (Caseys *et al*. 2021; Pink *et al*. 2022; Derbyshire *et al*. 2022). Thus, it remains unclear if the changes noted in large-effect co-transcriptome studies are transposable to a system in which numerous signals are varying in both the host and pathogen. To decipher the influence of regulatory variation in stem rust resistance, a host focused transcriptome study on barley (*Hordeum vulgare*), showed that host transcripts are largely controlled by a plethora of quantitative moderate effect loci involving a diversity of mechanisms and pathways (Druka *et al*. 2008; Moscou *et al*. 2011). A transcriptomic study on strains of the wheat pathogen *Zymoseptoria tritici* differing in virulence, found conserved and non-conserved gene expression patterns in genes involved in virulence, suggesting that heterogeneity in pathogen transcriptome contributes to quantitative virulence (Palma-Guerrero *et al*. 2017). This suggested that at least in the pathogen, quantitative virulence is linked to quantitative variation in the transcriptome.

However, it is unclear how the bidirectional nature of a host-pathogen interaction responds to quantitative variation in the pathogen. A co-transcriptome approach is required to query how quantitative genetic variation in generalist quantitative host-organisms systems transmits between the two organisms via transcriptome variation ultimately leading to the phenotypic outcome (Corwin *et al*. 2016b; Soltis *et al*. 2020). In such cases, measuring the dual transcriptome of interacting partners simultaneously across multiple genotypes and constructing a dual transcriptomic network would aid in understanding network-for-network interaction, the flow of information happening at the transcriptome level (Zhang *et al*. 2017, 2019). While correlation does not capture the directionality and causality between variability at genome and transcriptome levels of the two species, integrating a genetic mapping approach can help decipher the direction of causality by which genetic variation in the host and pathogen influence the flow of information (Chen *et al*. 2010). Ultimately this may enable a more complete model as to how the interaction leads to a specific disease phenotype (Chen *et al*. 2010; Christie *et al*. 2017; Almeida-Silva and Venancio 2021).

To explore how quantitative genetics shapes the bidirectional flow of information, we conducted a co-transcriptomic genome wide association study of the *Botrytis cinerea*-*Arabidopsis thaliana* pathosystem. Botrytis is a necrotrophic fungal pathogen infecting a wide range of plants (>1400 species) including *A. thaliana* (Leisen *et al*. 2022). Botrytis is a highly polymorphic species with a wide range of virulence on different hosts and an extensive collection of single nucleotide polymorphisms (SNP) enabling GWAS studies (Rowe and Kliebenstein 2007; Williamson *et al*. 2007; Amselem *et al*. 2011; Staats and van Kan 2012; Atwell *et al*. 2015; Corwin *et al*. 2016a). Virulence is mediated by a complex set of mechanisms including the secretion of a cocktail of proteins, which includes several cell wall degrading enzymes, cell death inducing proteins (CDIPs); necrosis and ethylene inducing proteins (NEPs), and metabolites like botrydial or botcinic acid. Further, Botrytis is also known to secrete a collection of sRNA molecules that can potentially target the expression of different host mechanisms (Choquer *et al*. 2021). This wide plethora of diverse yet redundant virulence mechanisms facilitates Botrytis infection on a wide range of plants while also allowing extensive genetic variation in individual mechanisms, e.g. the botrydial and botcinic acid pathways have presence/absence variation (Siewers *et al*. 2005; Pinedo *et al*. 2008; Plesken *et al*. 2021). All these clearly suggest the presence of variability in genome-transcriptome-metabolome mediated signalling processes in Botrytis (Leisen *et al*. 2022). Thus, a collection of Botrytis isolates acts as an assemblage of pathogens that is each sending different information into the host to create different perturbations of the host-pathogen information flow. Combining co-transcriptomics with genetic diversity in the host and pathogen diversity in this system can help to illustrate how the host-pathogen transcriptomes respond to the variation in a quantitative interaction.

To identify the pathogen loci that can shape a co-transcriptome response, we conducted a comparative expression Genome wide association (eGWA) study using 96 different wildtype *B. cinerea* strains. These were infected on three different hosts, wildtype *A. thaliana* Columbia 0 (Col-0) and two Arabidopsis mutants deficient in major defense pathways, *coi1-1* (jasmonate insensitive) and *npr1-1* (deficient in SA-mediated defenses) to test how the hosts variation may shape the pathogens response (Soltis *et al*. 2020). This pathosystem has no identified large-effect loci and allows us to investigate network-for-network interactions that may be masked by large effect ‘gene for gene’ relationships. This allowed us to compare two contrasting models; the host-pathogen co-transcriptome could be largely shaped by loci within Botrytis acting either dependently or independently of the host genotype. To test between these models, we mapped Botrytis loci that influence variation in the Botrytis- Arabidopsis co-transcriptome using genome wide efficient mixed model association and further assessed the results using network ANOVAs, to test the quantitative or qualitative nature of gene expression hotspots. Our analysis demonstrates that the major hotspots in the pathogen transcriptome do link to causing hotspots in the host transcriptome, suggesting that global shifts in the pathogen are not responsible for the major host responses. Network ANOVA models showed that the pathogen responds specifically and largely quantitatively to host genotypes and not qualitatively even though the host genotypes used are qualitative mutants in major signaling pathways. Finally, we could identify instances of host-genotype specific epistatic interactions. Our study thus sheds light on the complex transcriptome-transcriptome interaction, happening at the host-pathogen interface and how it is modulated by the genetic diversity in the host and the pathogen.

## Results

### Genetic variability in the pathogen differentially modulates the host and pathogen transcriptomes

To understand the relative impact of genetic variation in the host and pathogen on the Botrytis-Arabidopsis co-transcriptome, we calculated each transcripts’ relative broad-sense heritability (H^2^) attributed to the hosts (host H^2^: Col-0, *coi1-1*, and *npr1- 1*) or the pathogens genetic variation (pathogen H^2^: genetic variation among 96 Botrytis strains). We also calculated the fraction of the total variance controlled by the interaction of the host and pathogens genetic variation (co-H^2^: Supplementary Table 1). All the Botrytis transcripts showed a similar behavior in predominantly being influenced by pathogen H^2^ and the interaction of host and pathogen, co-H^2^ (Fig. 1: Host H^2^_avg_:0.01, Pathogen H^2^_avg_:0.15, co-H^2^_avg_:0.12). Thus, even knockout mutations in the hosts SA/JA-signaling pathways do not have consistent effect across all pathogen genotypes but instead the host’s effect on the pathogen depends on the pathogen genotype (Fig. 1A) (Zhang *et al*. 2019).

**Fig. 1.**
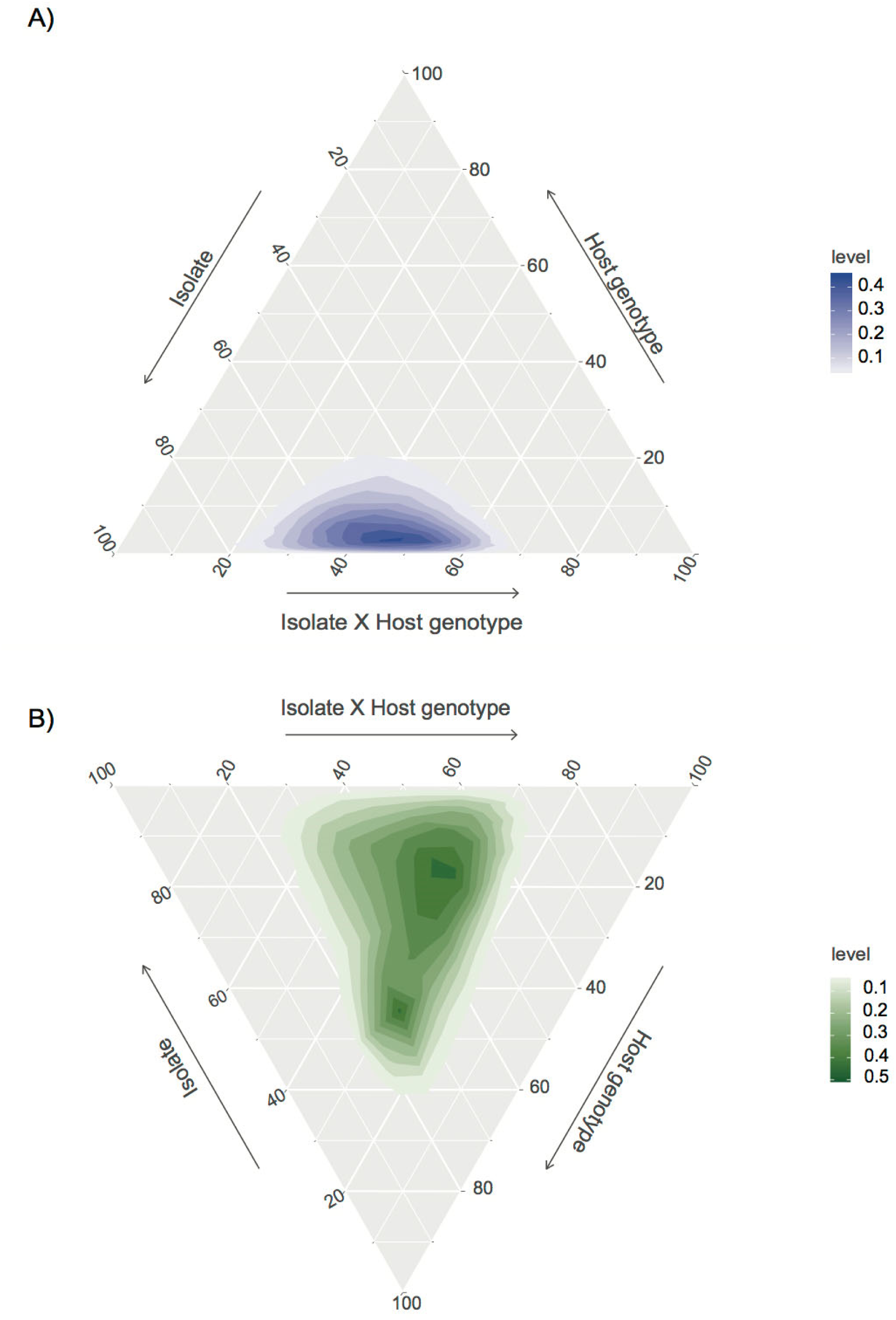
Differential genetic contributions to co-transcriptome variation. Shown are ternary plots representing the percentage of the total heritability that is attributable to the host, pathogen and host-pathogen interaction (different axes as labeled) on A) Botrytis transcripts B) Arabidopsis transcripts. The percentage of total heritability was determined by summing up the heritability attributed to host, pathogen and host-pathogen interaction terms and then dividing each individua term by that total.

The Arabidopsis transcripts showed a different pattern to the Botrytis transcripts with a wider spread dominated by a bimodal distribution (Supplementary Table 1 and Fig. 1B). One modality is near the center of the equilateral triangle where the Arabidopsis transcripts are equally influenced by host genotype, pathogen genotype and their interaction. This suggests that this modality has a set of host genes whose response to the pathogen is dependent on both the pathogen and the internal JA/SA pathway. The second modality is at a position where the transcripts had a nearly equal contribution of the pathogen and the host-pathogen interaction with little main effect from host genotype. These host transcripts would rely on the internal JA/SA signaling pathway in a manner that is completely condition on the pathogen’s genetic variation. These results imply that JA/SA signaling in the host is highly conditional on the pathogens genotype.

### Genomic distribution of co-transcriptome eQTLs

The above results show that the heritable genetic variation amongst the Botrytis strains influences the co-transcriptome via an interaction with the host genotype. This Host-Pathogen interaction could be caused by loci within Botrytis influencing the co-transcriptome and the identity of these loci may differ depending on the host genotype, i.e. wild-type (Col-0) specific or *coi1* specific pathogen loci. Alternatively, the causal loci in Botrytis may have a quantitative host conditionality whereby the same loci have effects in all host genotypes but the effect size changes depending on the host genotype. To test between these models, we mapped Botrytis loci that influence variation in the Botrytis-Arabidopsis co-transcriptome and assess how these pathogen loci are influenced by the host genotype.

To identify eQTL, we performed genome-wide association study across all detected Botrytis and Arabidopsis transcripts as measured separately on three different Arabidopsis genotypes (*coi1,* Col-0 *and npr1*). For the GWA we used the z-scaled expression values of 9,267 Botrytis genes and 23,947 Arabidopsis genes. For the Botrytis genetic polymorphisms, we used a previously generated dataset of Botrytis SNPs dataset consisting of 237,878 SNPs with a conservative minimum minor allele frequency cutoff of 0.20 (Soltis *et al*. 2019). eQTL mapping was conducted using Genome-wide Efficient Mixed Model Association (GEMMA) based on a univariate linear mixed model and a kinship matrix to account for the low but present population structure within the Botrytis collection. GWA was run separately on each transcript independently for each Arabidopsis-genotype. Previous work showed that given the large number of tests using the top SNP per transcript was an optimal compromise in minimizing the potential for false positives while maximizing the information available to identify genomic patterns for this analysis (Soltis *et al*. 2020). Thus, for further analysis we focused only on the most significant SNP per transcript.

Using these results, we first queried the genomic distribution of loci associated with variation in the Botrytis transcriptome. SNPs influencing a transcripts abundance can be located within the gene causing a direct effect such as altering the promoter, cis, or they can be located distal to the gene and alter the regulatory or other machinery influencing the gene, trans. To get an overview of the distribution of eQTLs in Botrytis, cis/trans plots (Fig. 2) were generated for the Botrytis transcripts separately, as measured on each Arabidopsis genotype. In these plots, the genomic position of the top SNP for each transcript (x-axis) is plotted against the genomic position of the gene encoding the transcript (y-axis). However, there is evidence for trans-eQTL hotspots, which can be seen as vertical lines of points. Trans-hotspots represent Botrytis polymorphisms that are associated with the variation in transcript abundance for a large number of transcripts and typically function in trans to the associated transcripts. Further, these hotspots differ across the three host genotypes using this approach (Fig. 2). As previously found, there is a paucity of cis-eQTL within this pathogen as indicated by the absence of a cis diagonal on any of the three host genotypes.

**Fig. 2.**
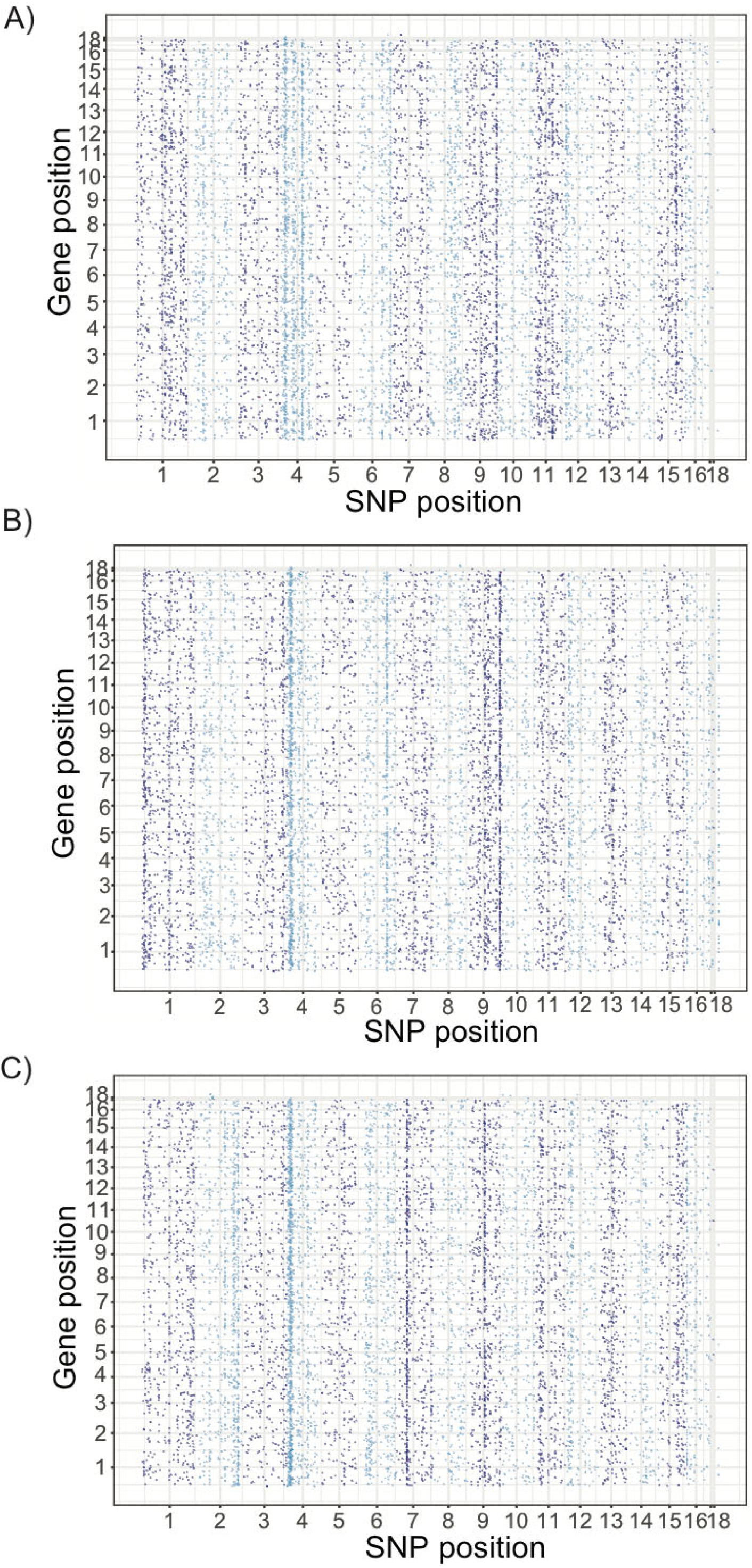
Distribution of SNPs associated with transcript variation in *Botrytis cinerea.* Comparison of the eQTL associated SNPs for each Botrytis transcripts in each Arabidopsis genotype A) *coi1* B) Col-0 and C) *npr1.* The position on the x-axis show the single most significant SNP found to affect a given Botrytis transcripts for which the genomic center along the 18 chromosomes is plotted on the y-axis. The positions of the SNPs have with two alternating colors to distinguish the chromosomes. Purple and blue are used to indicate the alternating chromosomes.

### Trans-eQTL hotspots vary across host genotypes

To investigate trans-hotspots, we considered the host genotype as an environment that plastically shapes the pathogen genotypes. To compare the hotspots, we plotted the number of Arabidopsis and Botrytis transcripts significantly associated with each SNP using only the top SNP per transcript for each host genotype for a total of six datasets (Fig. 3). Using random permutations, the thresholds for a hotspot were, ≥ 20 Botrytis transcripts per SNP and ≥100 Arabidopsis transcripts per SNP were used (Soltis *et al*. 2019). Studying the three different host genotypes revealed a potential major effect of the host has on the co-transcriptome. Changing the host genotype altered the number of hotspots with 26 eQTL hotspots for Botrytis transcripts when infecting *coi1* and 18 on *npr1,* while 22 eQTL hotspots were detected in Arabidopsis wild type host, Col-0 (Supplementary Table 2). Additionally, the majority of the eQTL hotspots for Botrytis transcripts (46 out of 55) were unique to individual Arabidopsis host genotypes with only two eQTL hotspots shared across all three host genotypes, four eQTL hotspots shared between *coi1* and Col-0, one between Col-0 and *npr1* and two between *npr1* and *coi1* (Fig. 4A).

**Fig. 3.**
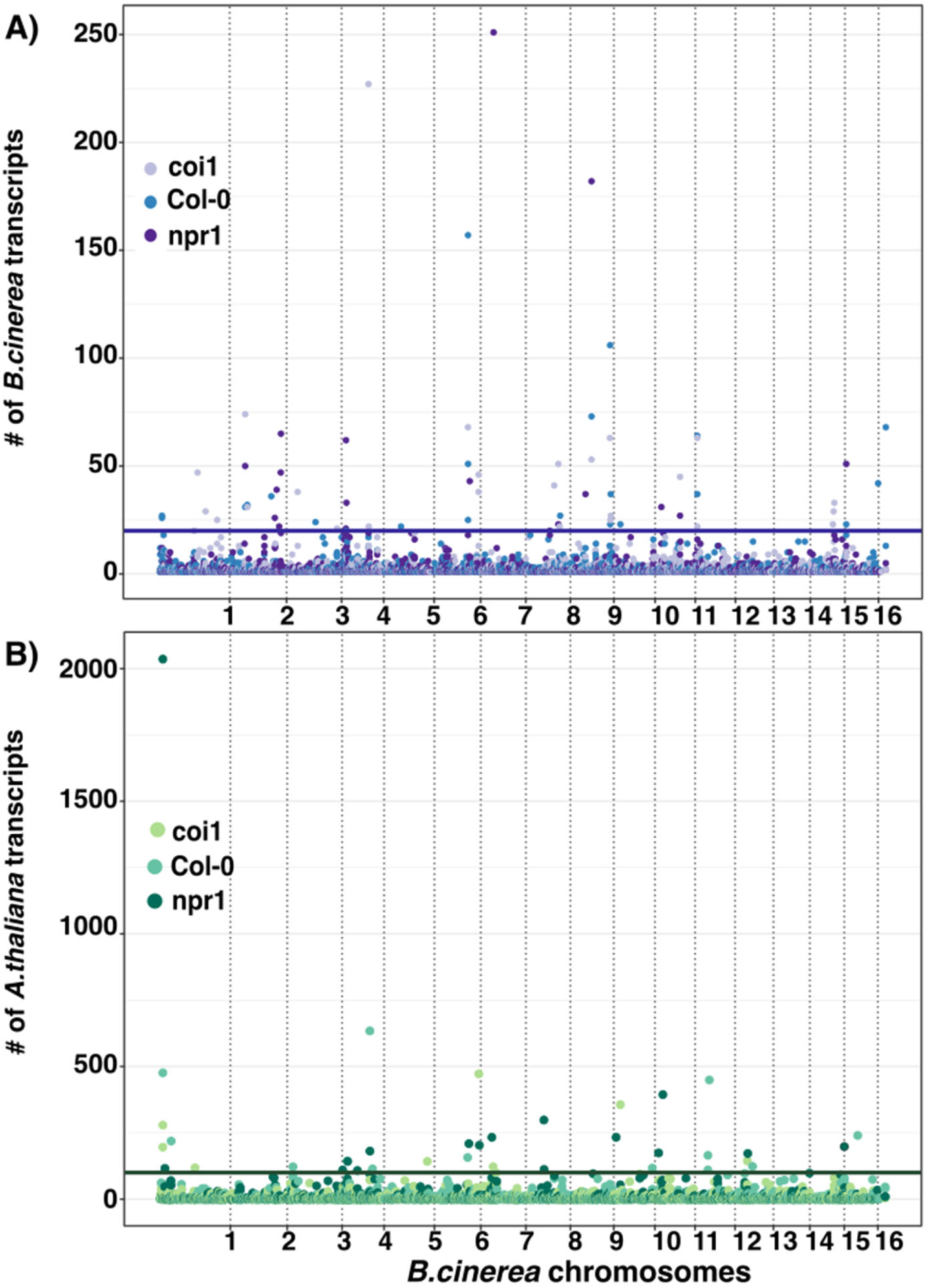
Distribution of Arabidopsis and Botrytis transcript/SNP associations across the Botrytis cinerea genome. Shown are Manhattan-like plots representing the number of genes associated with a specific eQTL. Hyphenated lines indicate the significant thresholds for a hotspot fixed based on permutation and randomization ≥ 20 transcript/ SNP (for Botrytis transcripts) and ≥100 transcripts/ SNP (for Arabidopsis transcripts) (Soltis *et al. 2020*), A) shows the number of Botrytis transcripts whose variation associates with each SNP when using the transcriptomes from isolates infected on Arabidopsis *coi1*, Col 0 and *npr1* per legend. B) shows the number of Arabidopsis transcripts whose variation associates with each Botrytis SNP when using the transcriptomes from isolates infected on Arabidopsis *coi1*, Col 0 and *npr1*.

**Fig 4.**
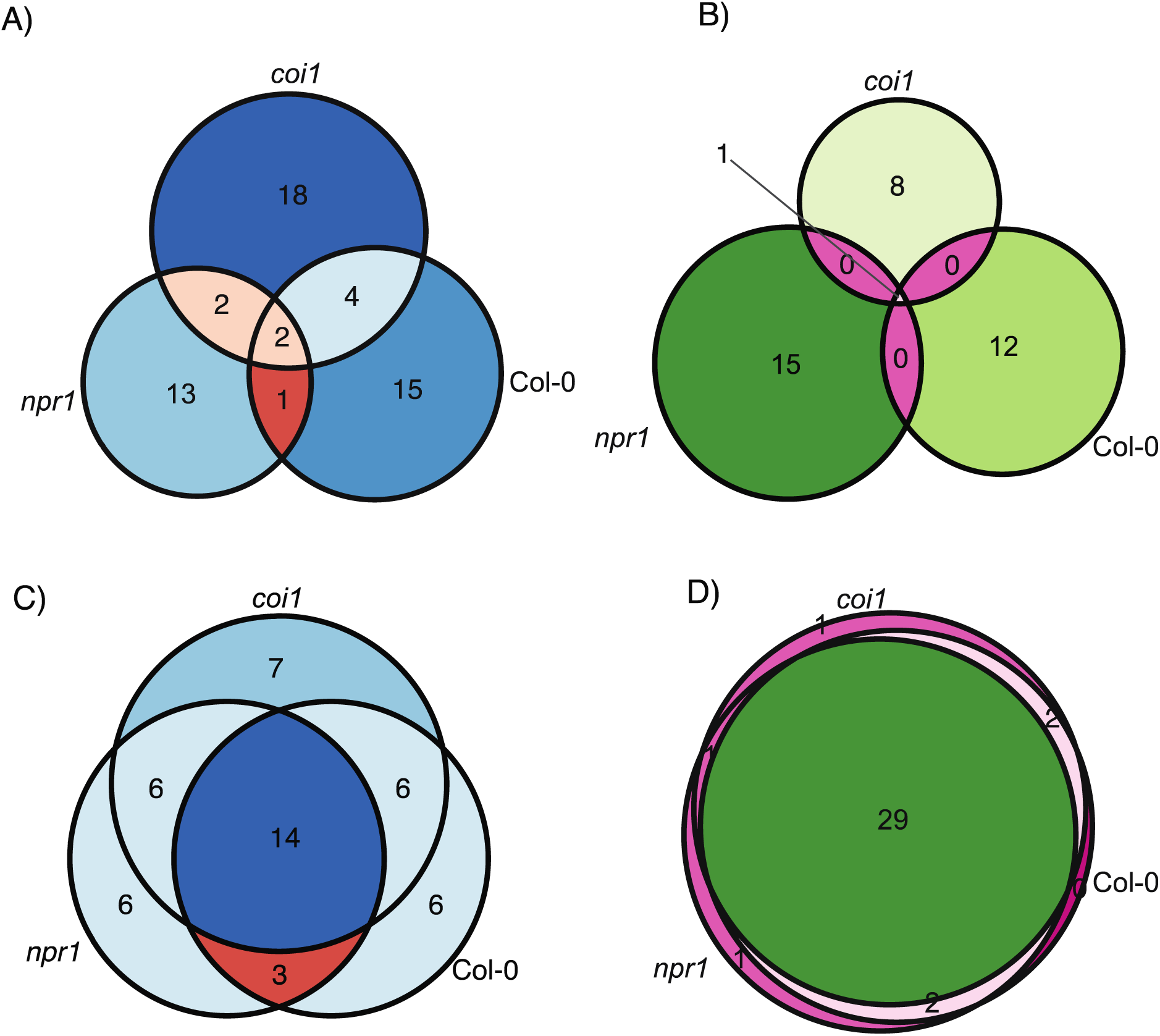
Effect of host genotype on eQTL hotspot identification. A) and B) present the number of GEMMA expression hotspots found using individual transcript GWA for Botrytis transcripts (hotspot n = 55) and Arabidopsis transcripts (hotspot n = 36) respectively and how they distribute across the three host genotypes. C) and D) present the number of gene expression hotspots identified using the network transcript-based approach to test each hotspot using a single host model of Botrytis transcripts (significant hotspot n = 48) and Arabidopsis transcripts (significant hotspot n = 36) respectively.

Shifting from the Botrytis transcriptome to the Arabidopsis transcriptome, found a similar pattern with 36 total eQTL hotspots, only one of which is identified across multiple host genotypes (Fig. 4B., Supplementary Table 3). Thus, the host genotype influences the ability to identify Botrytis SNPs that link to trans eQTL hotspots in the co-transcriptome. We next tested if any hotspots were shared across the two species transcriptomes as would be expected if a Botrytis SNP influences the Botrytis transcriptome consequently altering the Arabidopsis transcriptome. Across all the host genotypes, there was only a single trans-eQTL hotspot identified as influencing the transcriptome of both pathogen and host transcriptomes (Supplementary Tables 2 and 3). This trans-eQTL hotspot was found in the *coi1* host genotype and the SNP is within the *Bcin06g07340* gene encoding a synonymous polymorphism in a FAD binding domain protein of unknown function. Future work is needed to ascertain if this is a causal association and what may be creating the lack of connectivity between hotspots in the two species.

### Single host modeling of eQTL hotspots

One difficulty with the eQTL hotspot approach is that it relies upon combining independent tests across transcripts that are each susceptible to stochastic noise. We thus proceeded to investigate these eQTL hotspots using network-based linear models (Network Model) to more directly assess the influence of the SNPs on sets transcripts (Kliebenstein *et al*. 2006). To implement the network approach, we defined a network as the transcript set linked to each specific eQTL hotspot and the set of transcripts were used in the model. Given the potential for genome structure or other data structure to influence the significance estimates, we generated empirical *p*-value distributions by permutation testing to empirically estimate the alpha error potential. For each network, 100 random sets of transcripts of the same membership size were generated, and the linear modeling was performed using these random transcript sets. This generated a random distribution of 100 models. This showed that there was some bias in the p-value distribution, due to genome, population or other data structure, and as such an ANOVA term was only considered significant if the *p*-value was ≤ 0.05 and it was within the 5% tail of empirical permutations (empirical a = 0.05).

The initial round of network models focused on each Botrytis trans-eQTL hotspot in single Arabidopsis genotypes (single host models). All 55 identified eQTL hotspots influencing the Botrytis transcriptome were tested on all three-host genotypes to test if the host genotype dependency might be an issue of GWA power. Using this single host model, seven of the 55 eQTL hotspots were not significant while 48 eQTL hotspots for Botrytis transcripts, were found to be significant in at least one of the three Arabidopsis genotypes (Fig. 4C, Supplementary Tables 2 and 4). Interestingly, this single host model showed that 29 of the eQTL hotspots were detected on multiple hosts. Thus, the percentage of Botrytis transcript eQTL hotspots detected on multiple hosts increased from 16% with the single transcript GWA to 40% with the Network Model. This increase included, 14 of the 48 eQTL hotspots for Botrytis transcripts being found on all the three-host genotypes. This suggests that Network Models are more sensitive in detecting the quantitative effects at eQTL hotspots.

Applying the single-host Network Model to eQTL hotspots for Arabidopsis transcripts showed that all 36 eQTL hotspots were found to be significant in at least one of the host genotypes (Supplementary Tables 2 and 4). Twenty-nine eQTL hotspots for Arabidopsis transcripts were common to all the three Arabidopsis genotypes, while a single eQTL hotspots was specific to each *coi1* and *npr1* (Fig. 4D). No eQTL hotspot was specific to the wild-type genotype Col-0. Consistent with results of the Network Model for Botrytis transcripts, the number of Arabidopsis eQTL hotpots significant on multiple hosts increased considerably, from 3% in individual transcript GWA to 94% when we used the Network Model (Supplementary Table 4). One interpretation of this result is that there isn’t absolute host specificity but possibly quantitative variation or plasticity of the transcriptomes in response to the different Arabidopsis host genotypes.

### Multi-host modeling of eQTL hotspots provides evidence of host genotype effect

To directly test for quantitative host by pathogen genetic interactions at the above eQTL hotspot loci, we combined the host genotypes into a multi-host network linear model. This multiple host network model specifically tests for the significance of SNP- Host genotype interactions across the set of transcripts influenced by the eQTL hotspot. We again utilized the permutation approach as described to estimate significance thresholds. Our focus was on the SNP and SNP by Host Genotype interaction terms within the model. The SNP term directly tests the main effect of the hotspot SNP on the transcripts across all three Arabidopsis genotypes, whereas the SNP by Host Genotype term tests if the SNP has an interaction effect i.e., the influence of the SNP on the transcript network differs across the host genotypes. To visualize the interaction of the host genotype with each SNP, allele specific average expression values heat-maps and line plots for network transcripts were generated, for the three Arabidopsis genotypes (Fig. 5, Fig 6). Isolates with null alleles were not included in the analysis.

**Fig 5.**
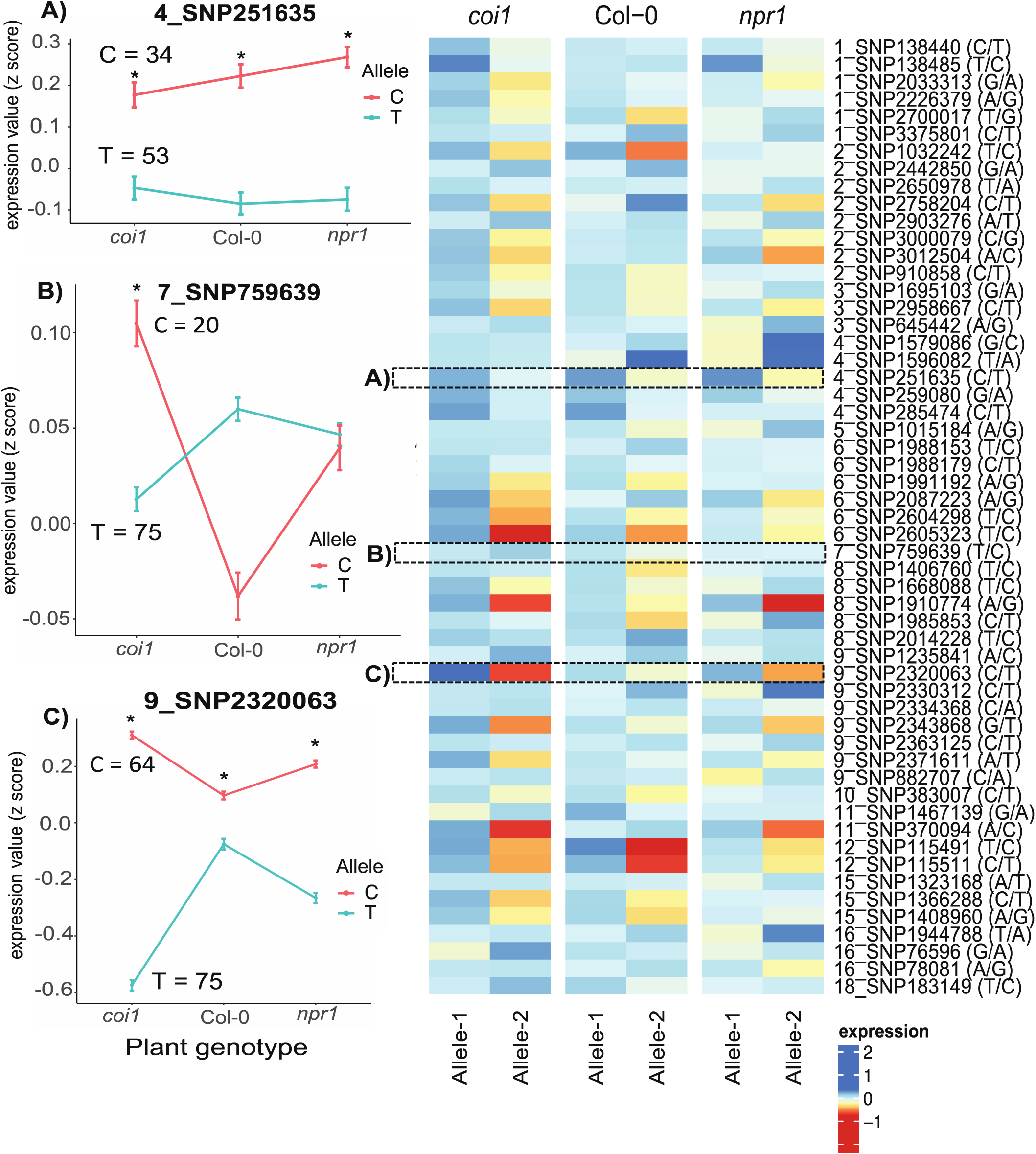
Estimating Botrytis trans-eQTL hotspot effects on Botrytis networks using network transcript z-scores. Estimated Botrytis SNP effects on Botrytis transcript networks as measured using averaged z-scores. For each trans-eQTL hotspot, the transcripts significantly associated with this SNP were grouped as a network and the average z-score across the network was used to estimate network expression. The results from all hotspots are shown in the heatmap with the SNP position indicated. A) example of a SNP with a significant host-genotype main effect and no host x pathogen genotype interaction effect B) example of a SNP with significant interaction of host x pathogen genotype but no pathogen main effect C) example of a SNP with a significant host-genotype main effect and a significant host x pathogen genotype interaction.

**Fig 6.**
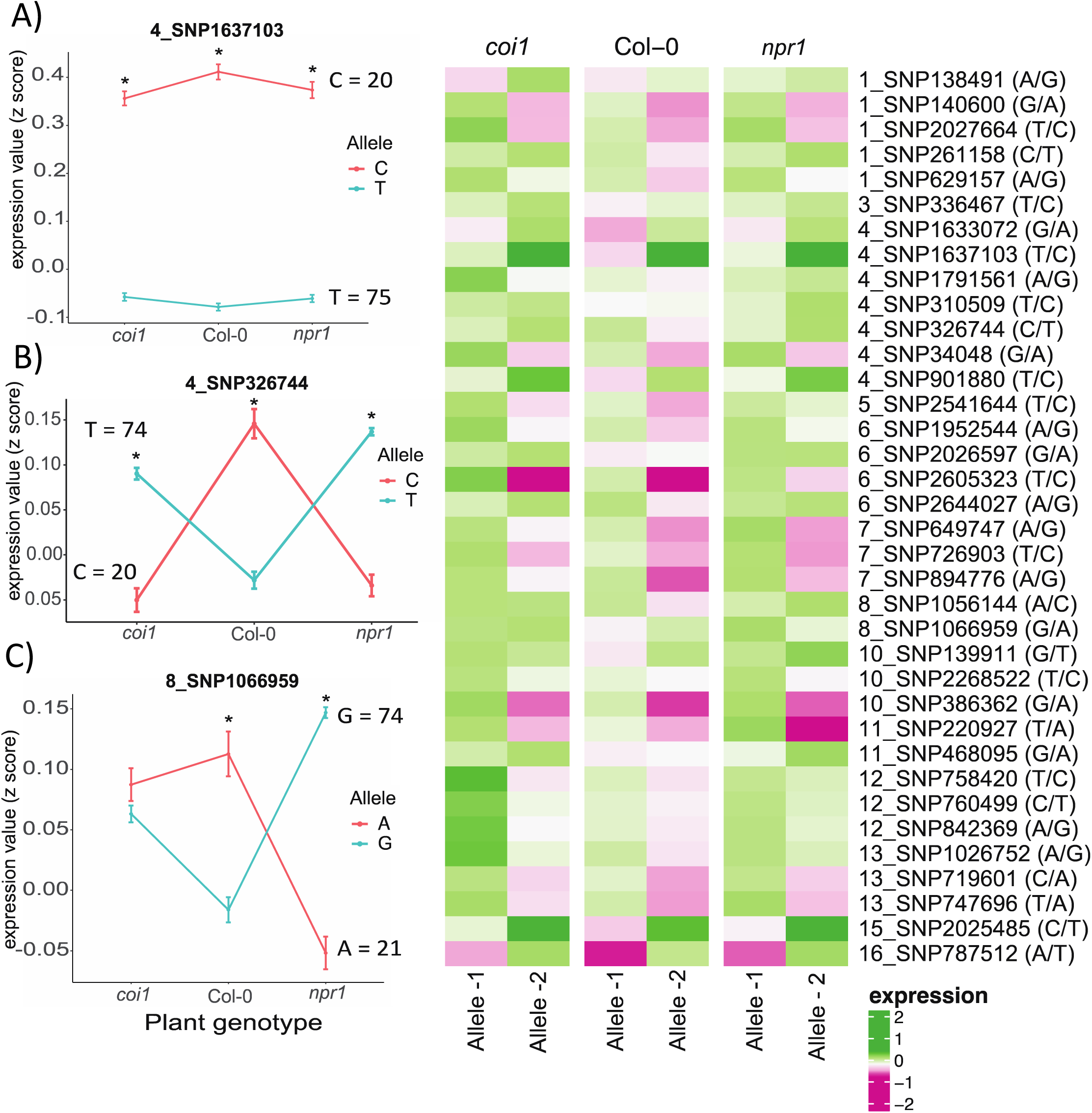
Estimating Botrytis trans-eQTL hotspot effects on Arabidopsis networks using network transcript z-scores. Estimated Botrytis SNP effects on Arabidopsis transcript networks as measured using averaged z-scores. For each trans-eQTL hotspot, the transcripts significantly associated with this SNP were grouped as a network and the average z-score across the network was used to estimate network expression. The results from all hotspots are shown in the heatmap with the SNP position indicated. A) example of a Botrytis SNP with a significant host-genotype main effect but no host x pathogen genotype interaction effect B) example of a SNP with significant interaction of host x pathogen genotype but no pathogen main effect C) example of a SNP with a significant host-genotype main effect and a significant host x pathogen genotype interaction.

Most eQTL hotspots had a significant main effect across the three host genotypes; 38 of 55 Botrytis eQTL hotspots and 32 of 36 Arabidopsis eQTL hotspots (Supplementary Tables 2 and 3). The seven eQTL hotspots in the Botrytis transcriptome found as not significant in the single host models remained non-significant in the multi-host model. 33 of the Botrytis eQTL hotspots and 24 of the Arabidopsis eQTL hotspots (based on network analysis) that had main effect on the respective transcripts also had significant interaction effects with the host genotypes, indicating that they affected the network transcripts significantly across all the Arabidopsis genotypes however their effect varied quantitatively across the different Arabidopsis genotypes (example: SNP: 9_SNP2320063; Fig 5C and example SNP: 4_SNP326744; Fig 6B). A further 13 Botrytis eQTL hotspots and four Arabidopsis eQTL hotspots were found to have solely host genotype specific effects (example: 7_SNP759639; Fig 5B; SNP: 8_SNP1066959, Fig 6C). Finally, a few eQTL hotspots, 5 Botrytis and 8 Arabidopsis, displayed only a main effect (e.g. solely pathogen genotype) with consistent effects across all host genotypes (example: 4_SNP251635; Fig 5A; 4_SNP1637103 Fig 6A). These results further suggest that most networks influenced by genetic variation in the co-transcriptome show a host x genotype related plasticity and this plasticity is largely quantitative in nature.

### Evidence for host genotype specific epistatic interactions

The above analysis suggested that a majority of eQTL hotspots for Botrytis and Arabidopsis transcripts were unrelated with only a single locus being a hotspot for both species’ transcriptomes. This suggested another hypothesis: Botrytis transcripts/loci influencing the Arabidopsis eQTL hotspots may be linked in trans to Botrytis eQTL hotspots. To test this possibility, we queried for Botrytis genes that has a SNP associated with an Arabidopsis eQTL hotspot. We then cross-referenced this list of Botrytis genes that may cause Arabidopsis transcript variation to test if these genes transcripts were controlled in trans by a Botrytis eQTL hotspot. This query identified five Botrytis genes with variation linked to Arabidopsis eQTL hotspots and their transcript variation is linked to 8 different eQTL hotspots for Botrytis transcripts (Supplementary table 6). This suggests that the Botrytis trans hotspot should work through the Botrytis gene associated to the Arabidopsis hotspot suggesting a possibility of epistasis between the 2 SNPs in Botrytis. To test if there was evidence for epistasic interactions between the two SNPs in modulating the Arabidopsis transcriptome we used Network Models. Here again both single host model and multiple host model was used. Using this approach showed that a majority of the 8 potential epistatic interactions were significant in *coi1*, Col-0 and *npr1* using the single host model. Further, using the multiple host model showed that six of the eight interactions were found to be significant across all the genotypes and also significant host genotype specific effect (Supplementary table 6). This suggests that it is possible in co-transcriptomics to use both the host and pathogen transcriptome to identify potential epistatic interactions wherein a Botrytis transcriptome hotspot influences a Botrytis transcript that is associated with an Arabidopsis transcriptome hotspot.

### Enrichment of enzymatic activities in genes containing trans-eQTL hotspot SNPs

To investigate the genes and possible polymorphisms underlying the identified eQTL hotspots we queried the annotation of the genes containing the SNP and the potential effect of the SNP on the genes function. Average linkage disequilibrium (LD) decay in the *B. cinerea* genome is < 1kb (Atwell *et al*. 2018), hence we focused on genes where the eQTL hotspot SNP was located plus or minus 1 kb of the start/stop codon. 36 out of the 55 eQTL hotspots for Botrytis transcripts and 28 out of 36 eQTL hotspots for Arabidopsis transcripts located within a gene and the rest were intergenic. Nine of the hotspots for Botrytis transcripts and 12 of the eQTL hotspots for Arabidopsis transcripts were linked to two adjacent genes. Using these gene lists, we queried if there was any enrichment in the potential function of these potential causal genes (Supplementary Table 7). This showed that for the genes underlying the Botrytis eQTL hotspots there was an enrichment for ubiquitin and enzymatic processes (Supplementary Table 7A ). The genes underlying the Arabidopsis eQTL hotspots showed enrichment for enzymatic processes especially ones in folate and sulfur metabolism (Supplementary Table 7B). Trans-eQTL hotspots are often thought to be linked to transcription factors but there was no enrichment for transcription factors in the genes containing SNPs linked to these trans-eQTL hotspots (Supplementary Table 7).

### Potential functions of transcript networks influenced by trans-eQTL hotspots

To better understand the potential networks modulated by the eQTL hotspots, we investigated the function of the transcripts linked to each of these eQTL hotspots. We first queried the Arabidopsis transcript networks using Gene ontology (GO) enrichment analysis for over-represented biological processes. As previously found, GO analysis revealed that eight of the hotspots for Arabidopsis transcript networks displayed an overrepresentation of photosynthesis-related functions (Zhang *et al*. 2017, 2019). Five of the hotspots were enriched in genes related to abiotic stress, 6 enriched in genes related to biotic stress and 1 of the gene clusters was enriched in genes involved in the metabolism of specialized metabolites, including glucosinolates. However, while these enrichments are known to be linked to host-pathogen interactions, they are fairly vague. To dive into more specific mechanism, we conducted network enrichment using specific networks previously linked to Botrytis resistance (Zhang *et al*. 2017, 2019). This showed that 10 of the 36 eQTL hotspots for Arabidopsis transcripts were enriched in genes belonging to Network 1 which consists of genes related to JA/SA signaling and the production of indolic phytoalexins known to defend against Botrytis (Fig. 7). Eleven of the eQTL hotspots for Arabidopsis transcripts were enriched in genes belonging to network 4 that are enriched in nuclear-encoded photosynthetic genes localized on chloroplasts (Fig. 7). Additionally, the trans-eQTL hotspot analysis identified a number of new networks that did not readily have GO or a priori identifiable function.

**Fig 7.**
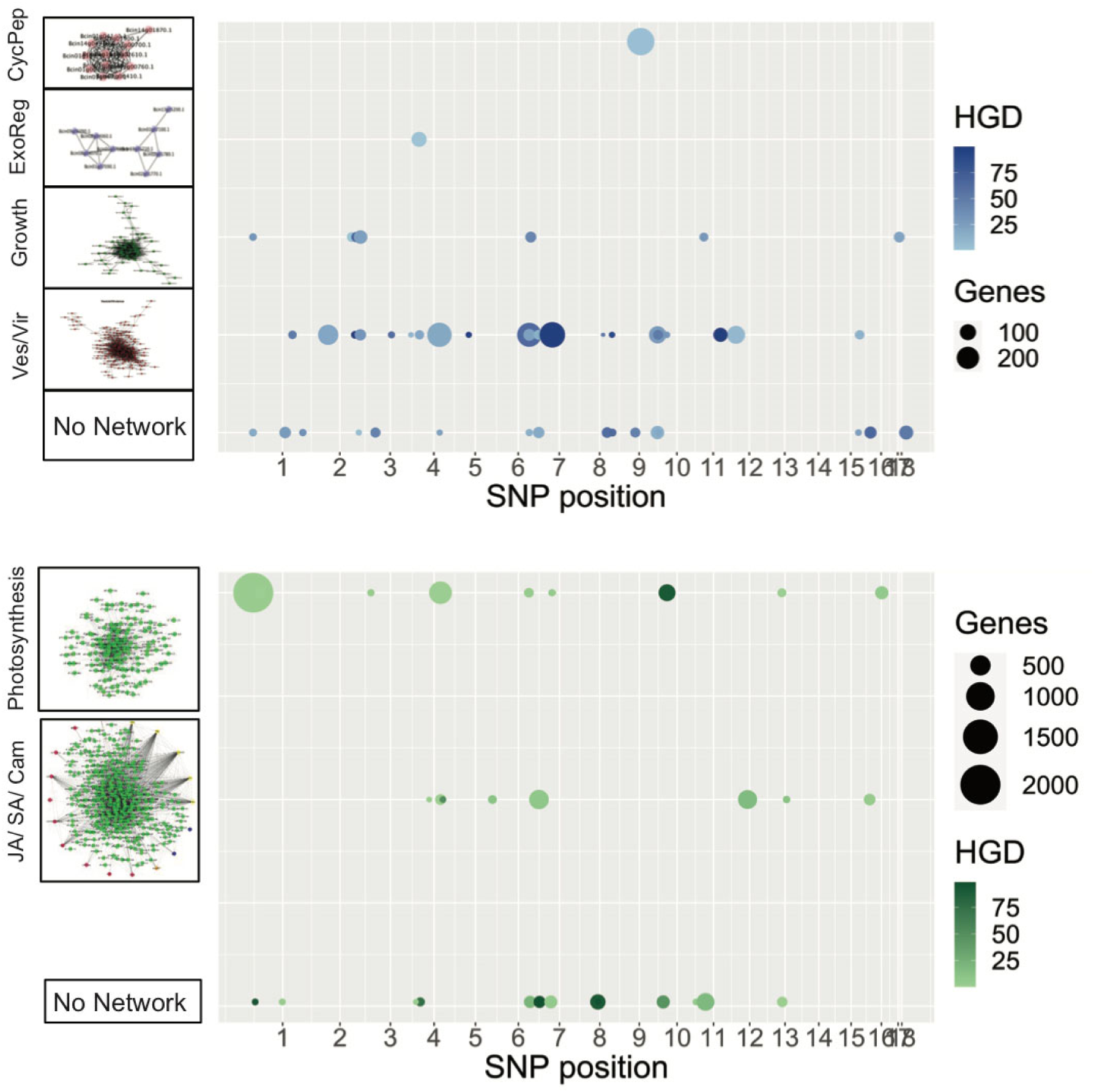
Polygenic manipulation of key virulence associated networks. The distribution of SNPs found to have an enriched association with previously identified transcript/biological networks and the fractions of the network influenced by the SNPs are shown. The position of the circles along the x axis show the position of the SNP and the radius of the circle is proportional to the number of genes within the network that are influenced by that SNP. A) trans-eQTL hotspots for Botrytis networks B) trans-eQTL hotspots for Arabidopsis networks. The network names are based on biological functions from gene ontology analysis of network members, from Fig. 4 of Zhang *et al*. (2019) and Fig. 6 of Zhang *et al*. (2017). Ves/Vir: vesicle/virulence network, Growth: Translation/growth network; ExoReg: exocytosis regulation network; CycPep: cyclic peptide network; JA/SA/Cam: JA and SA signaling processes and camalexin biosynthesis network. A Hypergeometric test was used to test for over-enrichment in genes targeted by each eQTL hotspot for genes found in the previous Botrytis and Arabidopsis transcriptome modules.

Because GO annotation of networks is limited in Botrytis we focused on using the same prior network analysis to query the potential function of the Botrytis transcript eQTL hotspots (Zhang *et al*. 2017, 2019). This showed that there was a single trans-eQTL hotspot that controlled all members of a single biosynthetic gene cluster predicted to make Cyclic Peptides that can be associated with virulence (Fig. 7). Most Botrytis trans-eQTL hotspots (25) were enriched in genes that co-express with each other and are associated with the formation and movement of vesicles potentially related to altering virulence. Eight of trans-eQTL hotspots all associate with genes linked to increased translation and potentially growth rate. Further, there was a number of novel networks identified using the trans-eQTL hotspots (Fig. 7). Thus, the previously identified networks are all modulated by multiple eQTLs in this system suggesting that the polygenic basis of this co-transcriptome interaction may filter through a few common networks.

## Discussion

Plant-pathogen interactions involves the bidirectional exchange of information between the two interacting organisms that alters the organism’s transcriptomes. In specialist pathogens these interactions, are largely determined by single/ few large effect genes in either/or both species. In contrast, generalist pathogens like Botrytis utilize an array of genes with quantitative effects. How this quantitative interaction alters the bidirectional exchange of information and the mutual transcriptome responses is unclear. Here we utilized a collection of 96 different Botrytis strains to study the bidirectional flow of information in plant-pathogen quantitative interactions and how the host and pathogen genotypes influence these interactions.

In this study using host genotypes that abolish the key SA and JA immune signaling pathways and a diverse collection of Botrytis genotypes, we found that pathogen transcripts are largely dependent on pathogen variation or its interaction with the host immune system (Fig. 1A). In contrast, the host transcriptome had two populations of transcripts (Fig. 1B). One population of host transcripts mirrored the pathogen by being largely dependent on the pathogen and pathogen x host interaction with little host effect. A second population of host transcripts showed a balanced contribution from host, pathogen and host x pathogen interactions. It is intriguing that the host genotype has the least influence on variability of both host and pathogen, despite the host’s genotypes having knockouts in major SA and JA defense signaling pathways. This implies that the influence of JA/SA pathway regulation on Arabidopsis transcripts is highly conditional on the pathogen genotype. Further, the SA and JA defense signaling pathways only influence the pathogen dependent on the pathogens genotype. Thus, there is a bidirectional flow of information in the Arabidopsis/Botrytis interaction with the pathogen having genetic variation in the ability to modulate the hosts JA/SA defense signaling pathways.

### Host effect on transcriptional plasticity

Transcriptional plasticity achieved by mutations in regulatory regions are known to be associated with many complex adaptive traits in several species including plant pathogenic fungi (Bódi *et al*. 2017; Krishnan *et al*. 2018; Haueisen *et al*. 2019). A recent study on *Fusarium virguliforme* a generalist pathogen suggest that it utilizes transcriptional plasticity to modulate infection strategies on wide range of morphologically and biochemically diverse hosts (Baetsen-Young *et al*. 2020). Similarly, in a generalist pathogen like Botrytis, transcriptional plasticity might be linked to an ability to sense the defense capability of the host. The transcriptional plasticity could facilitate optimal host invasion and adaption to numerous hosts, thus contributing to rapid evolution (Frantzeskakis *et al*. 2020). In our study, the differences in effects across host genotypes are a direct measure of host-modulated plasticity. Therefore, host modulated plasticity is a dominant component influencing both of the transcriptomes (Figs. 1-7).

Plasticity could be qualitative in nature such that an eQTL was identified on only one host genotype and no other suggesting that the eQTL may influence a function optimized to that one host. Alternatively, the plasticity could be quantitative whereby the eQTL influences the co-transcriptome across all or most host genotypes with a differing range of effects. In both the Botrytis and Arabidopsis transcriptomes, there was exclusively quantitative plasticity whereby the eQTL effects were present in each host albeit with different effects (Fig. 5 and 6). Further, the networks influenced by the plasticity were almost entirely controlled by a polygenic architecture such that each transcriptome network was linked to multiple eQTL (Fig. 7). Thus, host-Botrytis interactions are likely highly dependent on plasticity whereby each isolate of Botrytis makes different transcriptome decisions based on the specific host with which it is interacting. Correspondingly, this transmits signals to the host leading to different transcriptomes.

### Trans-eQTL hotspot causality

From the eQTL analysis, we were able to identify a large number of trans-eQTL hotspots controlling the co-transcriptome for both Botrytis (55) and Arabidopsis transcripts (36) (Fig. 3, 4 and 7). Trans-eQTL hotspots are a common feature of eQTL studies in both structured and unstructured populations. Frequently they are theorized to be major regulatory loci influencing a wide array of transcripts and this is frequently short-handed to mean that they are more likely to be transcription factors (Hansen *et al*. 2008).

While several studies have reported eQTLs in plant and pathogen genomes (West *et al*. 2007; Chen *et al*. 2010; Christie *et al*. 2017; Wilkerson *et al*. 2022), it has not yet been widely determined if these loci are enriched for regulatory genes like transcription factors. Interestingly, in this analysis we did not find any significant GO enrichment for transcription factors in the genes underlying the trans-eQTL hotspots. Other studies have found a similar paucity of transcription factors in eQTL studies (Weiser *et al*. 2014; Wang *et al*. 2018). In contrast, we did find GO enrichment for enzymatic functions underlying these trans-eQTL hotspots. This is not unprecedented as Arabidopsis trans-eQTL hotspots have been causally linked to both genes in primary and specialized metabolism (Kerwin *et al*. 2011; Francisco *et al*. 2021). Similarly, several studies also showed an enrichment of genes involved in specialized metabolism among the genes underlying trans-eQTL hotspots (Weiser *et al*. 2014; Wang *et al*. 2018). Thus, it is possible that genetic variation in the plasticity of Botrytis-host interactions is being predominantly modulated by variation in enzymatic/metabolic processes. None of the genes underlying these loci have been previously associated with plant-pathogen interactions providing a rich source of candidate genes to pursue in the future.

This work shows the potential for co-transcriptome analysis to show how plastic the host and pathogen transcriptomes are in response to genetic variation in each other. Highly plastic transcriptome responses indicate that both the host and pathogen carefully shape their regulatory response to the blend of signals moving back and forth between the two interacting organisms. It remains to be tested if the plastic response leads to the optimal transcriptome for the interaction of if the plasticity instead creates a blend of beneficial and harmful transcriptome responses. This will require mutating the different outputs of the co-transcriptome and measuring the virulence consequence across an array of interactions. Understanding if and how plasticity may help to shape specific responses is key to engineering resistance in the future.

## Materials and Methods

### Transcriptome data used in the study

In this study, RNA-seq was used to quantify the expression of both Arabidopsis and Botrytis genes in Arabidopsis leaves infected with 96 different Botrytis strains independently. We retrieved expression profiles for all Botrytis and Arabidopsis genes from an earlier RNA-seq experiment (Zhang *et al*. 2017, 2019). Briefly, the RNA-seq data comprises of the gene expression values of Arabidopsis and Botrytis genes during interaction of three *A. thaliana* genotypes (Col-0, *coi1-1*, *npr1-1*) with a global collection of 96 different *B. cinerea* strains collected as single spores from natural infections of fruits and vegetable tissues (Zhang *et al*. 2017, 2019; Atwell *et al*. 2018; Caseys *et al*. 2021). The data was generated using four replicates in a randomized block design divided across two independent balanced experiments for all interactions. Fully mature and expanded Arabidopsis leaves were harvested five weeks after sowing and inoculated with 40 spores with one of 96 Botrytis strains, in a detached leaf assay (Denby *et al*. 2004; Corwin *et al*. 2016a; Zhang *et al*. 2017, 2019). Whole leaves were sampled at 16 hours post inoculation, for RNA isolation. RNA-seq libraries were generated following (Kumar *et al*. 2012) and RNA sequencing was performed on an Illumina HiSeq 2500 using single end reads 50 bp at the U.C. Davis Genome Center-DNA Technologies Core. RNA-seq reads were trimmed using the fastx toolkit (http://hannonlab.cshl.edu/fastx_toolkit/ commandline.html) and aligned to both the *A. thaliana* TAIR10.25 and *B. cinerea* B05.10 ASM83294v1 cDNA reference genomes. Gene counts were pulled from the resulting sam file using a combination of SAMtools (Langmead *et al*. 2009; Li *et al*. 2009; Staats and van Kan 2012) and custom R scripts, summed across gene models and normalized (Langmead *et al*. 2009; Li *et al*. 2009; Staats and van Kan 2012). TMM method was used for normalization of gene counts using the function calcNormFactors() from the “edgeR” package (Robinson and Smyth 2007; Robinson and Oshlack 2010; Bullard *et al*. 2010; Nikolayeva and Robinson 2014). The linear model was applied on the TMM normalized gene counts using function glm.nb() from the “MASS” package (Venables and Ripley, 2002). Model-corrected means and standard errors for each transcript along with variance components were estimated using a general linear model that assumed a gaussian distribution and the lsmeans V2.19 package (Lenth 2016; Zhang *et al*. 2017). Broad-sense heritability (H^2^) of each transcript was calculated as the proportion of variance due to the genetic variability in Botrytis strains, Arabidopsis genotype, or their interaction effects.

### Genome wide association mapping

GWA of both Botrytis and Arabidopsis transcripts were performed as described in (Soltis *et al*. 2020). A total of 9,267 B. *cinerea* gene expression values and 23,947 *A. thaliana* gene expression values across different genotypes of Arabidopsis infected with 96 strains of Botrytis. Briefly, z-scaled model-adjusted least square means of normalized gene counts of both the *A. thaliana* and *B. cinerea* transcripts (Zhang *et al*. 2017, 2019) were used as the phenotype for GWA. A total of 237,878 SNPs across 96 different botrytis strains mapped to the *B. cinerea* B05.10 ASM83294v1 genome (Atwell et al., 2018), were used for the association study. GWA was performed using GEMMA (Zhou and Stephens 2012) which follows a univariate linear mixed model. A standardized relatedness matrix was calculated in GEMMA to account for the population structure among Botrytis strains. GWA was performed separately for each Arabidopsis genotype in the study.

### Defining eQTL Hotspots

For defining eQTL Hotspots we considered only the top SNP associated with each in Botrytis and Arabidopsis transcripts as previously described (Soltis *et al*. 2020) This provides a relatively conservative approach that while allowing some false positives has been shown to provide useful information about the genome-wide pattern of associations.

Thus, we considered 9,267 SNP associated for the 9,267 *B. cinerea* transcripts and 23,947 SNP associated for the 23,947 *A. thaliana* transcripts for each Arabidopsis genotype. When identifying hotspots, defined as a SNP (Top 1 SNP) which is associated with multiple transcripts, we used a permutation approach to identify a conservative threshold. A random permutation threshold using 1000 permutations found the largest random hotspot to be 11 transcripts for Botrytis and 80 transcripts for Arabidopsis (Soltis *et al*. 2020). Thus, we defined eQTL hotspots as those Top 1 SNPs that are associated with 20 or more Botrytis transcripts or with 100 or more Arabidopsis transcripts.

### Validation and annotation of gene expression hotspots

Z-scaled (for each gene independently across strains) model-adjusted least square means of normalized gene counts of both the *A. thaliana* and *B. cinerea* transcripts were used for this study. Firstly, a single host Network Model was used to validate the gene expression hotspots. In this model, all of the transcripts associated with a trans-eQTL hotspot are utilized within the same model to maximize the ability to look at coordinated effects. A Network model was performed on the data from expression data from each genotype separately.

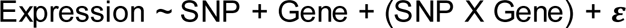

The main effects indicate the two alleles of the trans-eQTL hotspot SNP being tested and Gene represents the different transcripts associated with the trans-eQTL hotspot. *P*-values for each term were extracted and significance of each term in contributing to the variability of expression was analyzed. To make sure that the *p*-values were truly significant and was not significant just by chance, random sets of genes, spanning the entire genome of Botrytis or Arabidopsis were generated, which could be potentially be regulated by the hotspots. The same ANOVA model was run on using 100 random sets of transcripts of the same number as the transcripts for the hotspot to calculate the empirical estimate for each term. Only those terms where the empirical estimates were ≥ 95 were considered to be significant.

Next a multiple host model ANOVA was used to validate the gene expression hotspots across all the three Arabidopsis genotypes and to figure out if the polymorphisms displayed and Arabidopsis genotype specific effect of the expression of genes. ANOVA was performed on the pooled expression data of Arabidopsis genotypes.

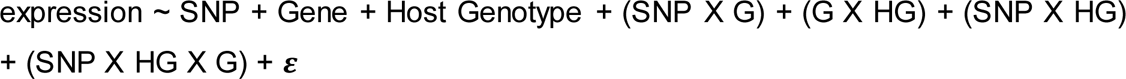

In this model expression denotes the expression value of each gene regulated by gene underlying the hotspot in all the holobionts, which includes all the three Arabidopsis genotypes, SNP denotes the different alleles underlying the trans-eQTL hotspot, Gene (G) denotes the individual transcripts associated with the trans-eQTL hotspot and Host Genotype (HG) is the three different Arabidopsis genotypes. *P*-values for each term were extracted and significance of each term in contributing to the variability of expression was analyzed. To make sure that the *p*-values were truly significant random sets of genes, spanning the entire genome of Botrytis or Arabidopsis were generated, which could be potentially be regulated by the hotspots. The same ANOVA model was run on 100 such random sets of genes to calculate the empirical estimate for each term. Only those terms where the empirical estimates were ≥ 95 were considered to be significant.

To further determine the functionality of each hotspot, we looked for the annotation of the genes underlying the hotspot. The SNPs were annotated with a gene by identifying if the SNP was within a distance of 1 kb upstream of the start codon of a gene or within 1 kb downstream of the stop codon of the gene This distance was chosen as the average linkage disequilibrium (LD) decay in the *B. cinerea* genome is < 1kb (Atwell *et al*., 2018). *B. cinerea* B05.10 ASM83294v1 GFF3 file was used to identify the genes underlying the hotspot, while gene functional annotations were obtained from the fungal genomic resource portal (fungidb.org). Further, SnpEff (Cingolani *et al*. 2012) was used to predict the effects of genetic variants underlying the hotspots.

### Epistasis

To test for the presence of epistasis, first we looked if any of the genes underlying Botrytis hotspots regulating Arabidopsis transcripts were present in the list of genes regulated by genes underlying Botrytis hotspots regulating Botrytis transcripts. Network ANOVAs were performed on such Botrytis genes which could possibly interact with each other and thus influence the Arabidopsis transcript, using single host model epistasis and multiple host model epistasis. For single host model epistasis the model was

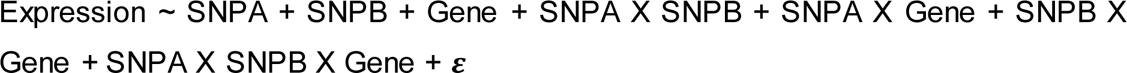

The main effects indicate the alleles of the two trans-eQTL hotspots, SNPA and SNPB, being tested and Gene represents the different transcripts associated with the trans-eQTL hotspot. *P*-values for each term were extracted and significance of each term in contributing to the variability of expression was analyzed. The same model was utilized for the multiple host epistasis by including a Host Genotype term and incorporating it into the various interaction terms. To make sure that the *p*-values were truly significant random sets of genes, spanning the entire genome of Botrytis or Arabidopsis were generated, which could potentially be regulated by the hotspots. The same ANOVA model was run on 100 such random sets of genes to calculate the empirical estimate for each term. Only those terms where the empirical estimates were ≥ 95 were considered to be significant.

### Enrichment analysis of the target gets

Gene ontology (GO) enrichment analysis for overrepresentation of molecular function & biological processes among the genes targeted by each eQTL hotspot in Arabidopsis was determined using the Bioconductor packages org.At.tair.db and topGO, R statistical environment. Hypergeometric test was conducted to look for over-enrichment in genes targeted by each eQTL hotspot for genes found in the previous *B. cinerea* and *A. thaliana* transcriptome modules (Subramanian *et al*., 2005; Zhang *et al*. 2017; Zhang *et al*. 2019).

### Gene co-expression analysis

To obtain genes co-expressed with a gene underlying a hotspot, we performed gene co-expression analysis. Z-scaled model-adjusted least square means of normalized gene counts of both the *A. thaliana* transcripts (23,947) and *B. cinerea* transcripts (9,267) from individual strain infection across three Arabidopsis genotypes were used. Spearman’s rank correlation coefficients of the gene expression values of the gene of interest with all other transcripts was calculated using the *cor* function in R. Three gene-for-gene correlation matrixes were generated independently for each of the three Arabidopsis genotypes. Transcripts which showed a correlation coefficient > 0.5 was considered co-expressed with the gene of interest.

### Statistics

All statistical analyses were performed in R environment using custom made scripts, including ANOVA, calculation of empirical estimates, GO enrichment analysis, and hypergeometric test

## Supplemental Tables

**Supplementary Table 1:** Estimated Broad-sense heritability (H2) of each transcript in Botrytis and Arabidopsis. Heritability refers to the fraction of total variance that is ascribed to the particular term.

**Supplementary Table 2.** List of GEMMA expression hotspots for Botrytis transcripts with the detailed annotation of the gene underlying the SNP and the number of transcripts in Botrytis regulated by the SNP.

**Supplementary Table 3.** List of GEMMA expression hotspots for Arabidopsis transcripts with the detailed annotation of the gene underlying the SNP and the number of transcripts in Arabidopsis regulated by the SNP.

**Supplementary Table 4:** Model estimated p-values and permutation derived alpha-error estimates from network ANOVAs.

**Supplementary Table 5:** Annotation of genes underlying hotspots for both Botrytis transcripts and Arabidopsis transcripts.

**Supplementary Table 6:** Summary of genes potentially under epistasis in Botrytis. True p-values and empirical estimates from network ANOVAs, estimated using single host epistasis model and multiple host epistasis model.

**Supplementary Table 7.** Summary of enrichment analysis of genes underlying hotspots for Botrytis and Arabidopsis transcripts.

